# STOPGAP, an open-source package for template matching, subtomogram alignment, and classification

**DOI:** 10.1101/2023.12.20.572665

**Authors:** William Wan, Sagar Khavnekar, Jonathan Wagner

## Abstract

Cryo-electron tomography (cryo-ET) enables molecular-resolution 3D imaging of complex biological specimens such as viral particles, cellular sections, and in some cases, whole cells. This enables the structural characterization of molecules in their near-native environments, without the need for purification or separation, thereby preserving biological information such as conformational states and spatial relationships between different molecular species. Subtomogram averaging is an image processing workflow that allows users to leverage cryo-ET data to identify and localize target molecules, determine high-resolution structures of repeating molecular species, and classifying different conformational states. Here we describe STOPGAP, an open-source package for subtomogram averaging designed to provide users with fine control over each of these steps. In providing detailed descriptions of the image processing algorithms that STOPGAP uses, we intend for this manuscript to also serve as a technical resource to users as well as further community-driven software development.

## Introduction

Cryo-electron microscopy (cryo-EM) combined with single particle analysis (SPA) has in recent years become a key method for determining the structures of biological macromolecules (Kühlbrandt, 2014). The ideal SPA specimens consist of a monolayer of purified molecules suspended in vitreous ice, with each molecule producing a randomly oriented projection in resultant electron micrographs. SPA determines structures by iterating between aligning each projection to a 3D reference structure and reconstructing an improved reference structure using the new alignment parameters. One limitation to SPA is that the specimen typically needs to consist of a monolayer of particles, otherwise the molecular projections begin to overlap one another in micrographs, thereby hindering alignment and structure determination.

There are a number of situations where overlapping structures cannot be avoided, including pleomorphic assemblies, membrane-associated complexes, and molecules in their near-native cellular environments (Beck & Baumeister, 2016). In these situations, key structural and biological information is inextricably tied to the complexity of the molecular environments, making the separation, isolation, or purification of molecular components undesirable. Cryo-electron tomography (cryo-ET) offers one approach to solving the problem of imaging overlapping molecules. In cryo-ET, rather than collecting a single images for each field a view, each field of view is imaged multiple times while tilting the specimen stage. This set of 2D projections, called a tilt-series, is then used to directly reconstruct a 3D representation of the field of view: a tomogram.

While tomograms overcome the overlap problem, they still suffer from a number of fundamental limitations. Cryo-ET specimens typically consist of either biological assemblies frozen into holey support grids, similar to single-particle cryo-EM specimens, vitrified cells, or cellular sections, such as focus ion beam (FIB) milled lamella. In each case, specimens have a roughly slab-like profile; when tilted, these slab-like specimens effectively become thicker with respect to the electron beam. This increasing thickness, coupled with physical restrictions of the microscope hardware, limits the angular range in which tomographic data can be collected. A common tilt range is from -60° to 60°, resulting in a specimen that is effectively twice as thick at the extreme tilt angles. Incomplete angular sampling, due to limitations in both the overall tilt range as well as the angular increments between tilt images, leads to unsampled or missing regions of Fourier space; this is referred to as the “missing wedge” (Schmid & Booth, 2008). While the missing wedge refers to unsampled regions of Fourier space, it also produces corresponding distortions in real space. Additionally, any other factors that affect electron micrographs also affect tomograms, such as the Contrast Transfer Function (CTF) (Wade, 1992) and electron exposure damage (Grant & Grigorieff, 2015), which fundamentally limits the amount of high-resolution information that can be obtained. As such, high-resolution structure determination from tomograms still requires the averaging of repeating structures across a dataset in order to fill the missing wedge, flatten CTF modulations, and increase the signal-to-noise ratio.

Determining structures from cryo-ET data requires a wide range of processing steps (Wan & Briggs, 2016). This includes steps to process raw 2D data into 3D reconstructions, such as tilt-series alignment, defocus determination, CTF correction, and tomographic reconstruction. The series of processing steps that starts after tomographic reconstruction and ends with the generation of higher-resolution averaged EM-density maps is analogous to SPA and typically referred to as subtomogram averaging (STA). STA includes a number of tasks such as 3D particle picking, often by template matching, the generation of higher-resolution EM-density maps by iterative subtomogram alignment and averaging, and subtomogram classification to separate heterogeneous structures.

Here, we describe STOPGAP, an open-source package for STA. STOPGAP uses a real-space correlation-based approach similar to a number of other STA packages (Förster *et al*., 2005; Hrabe *et al*., 2012; Nicastro *et al*., 2006; Castaño-Díez *et al*., 2012), but contains a several new algorithms that we describe here. At the core of many of STOPGAP’s algorithms it its treatment of the missing wedge, which improves the performance of template matching and subtomogram alignment, as well as the quality of averaged EM-density maps. We also describe algorithms for subtomogram classification by multireference alignment (MRA) that use stochastic approaches which allow for the assessing the reproducibility of classification results.

## Results

### Missing Wedge Model

Distortions caused by the missing wedge effectively result in structural features that are not present in the specimen or the isotropically-resolved reference maps produced in STA. As such, real-space comparisons between anisotropic tomographic data and isotropic references, can lead to poor results such as imprecise subtomogram alignment. To overcome this, real-space correlation STA packages apply a missing wedge filter to references prior to computing the cross-correlation with tomographic data; this is referred to as the constrained cross-correlation (CCC) (Förster *et al*., 2005; Frangakis *et al*., 2002; Schmid & Booth, 2008; Bartesaghi *et al*., 2008). The missing wedge filter is a Fourier-space filter that is meant to mimic the effects of tilt-series collection and tomographic reconstruction, i.e. the anisotropic sampling, in order to reproduce the real-space tomographic distortions into the isotropically-resolved reference. In real-space STA packages that use CCCs, the missing wedge filter is typically generated as a binary filter that includes all information between the maximum and minimum tilt angles (Förster *et al*., 2005; Hrabe *et al*., 2012; Nicastro *et al*., 2006; Castaño-Díez *et al*., 2012). While this accounts for the missing region caused by limited tilt range, it does not accurately describe the distortions in a tomographic reconstruction resulting from discrete angular sampling, CTF, and electron exposure damage.

In STOPGAP, the missing wedge filter is modelled to reflect the sampling geometry and amplitude modulations present in the tomographic data. This includes using Fourier slices rather than a solid wedge, CTF modulations, and exposure filtering. STOPGAP requires CTF-correction prior to or during reconstruction using methods such as tilted CTF correction (Xiong *et al*., 2009) or 3D CTF-correction (Turoňová *et al*., 2017; Kunz & Frangakis, 2017). STOPGAP also accounts for aliasing in the amplitude spectrum, which when combined with CTF-correction prior to or during tomogram reconstruction, allows for tight cropping of subtomograms to minimize computational costs while avoiding potential signal loss due to signal delocalization outside the subtomogram edges.

To illustrate the impact of different missing wedge components on the CCC, simulated 2D examples are shown in Figure 1. These examples represent tomographic planes orthogonal to the tilt axis, i.e. the standard XZ-plane in tomograms. Applying no missing wedge mask produces a peak in the cross-correlation map, but with significant background. Applying a continuous wedge or a per-tilt slice wedge produces a slightly sharper peak but still shows significant background correlation. While the per-tilt slice wedge has very similar performance to the continuous wedge mask, the slices also allow for applying tilt-image specific amplitude modulating factors such as CTF and exposure filtering.

**Figure 1:**
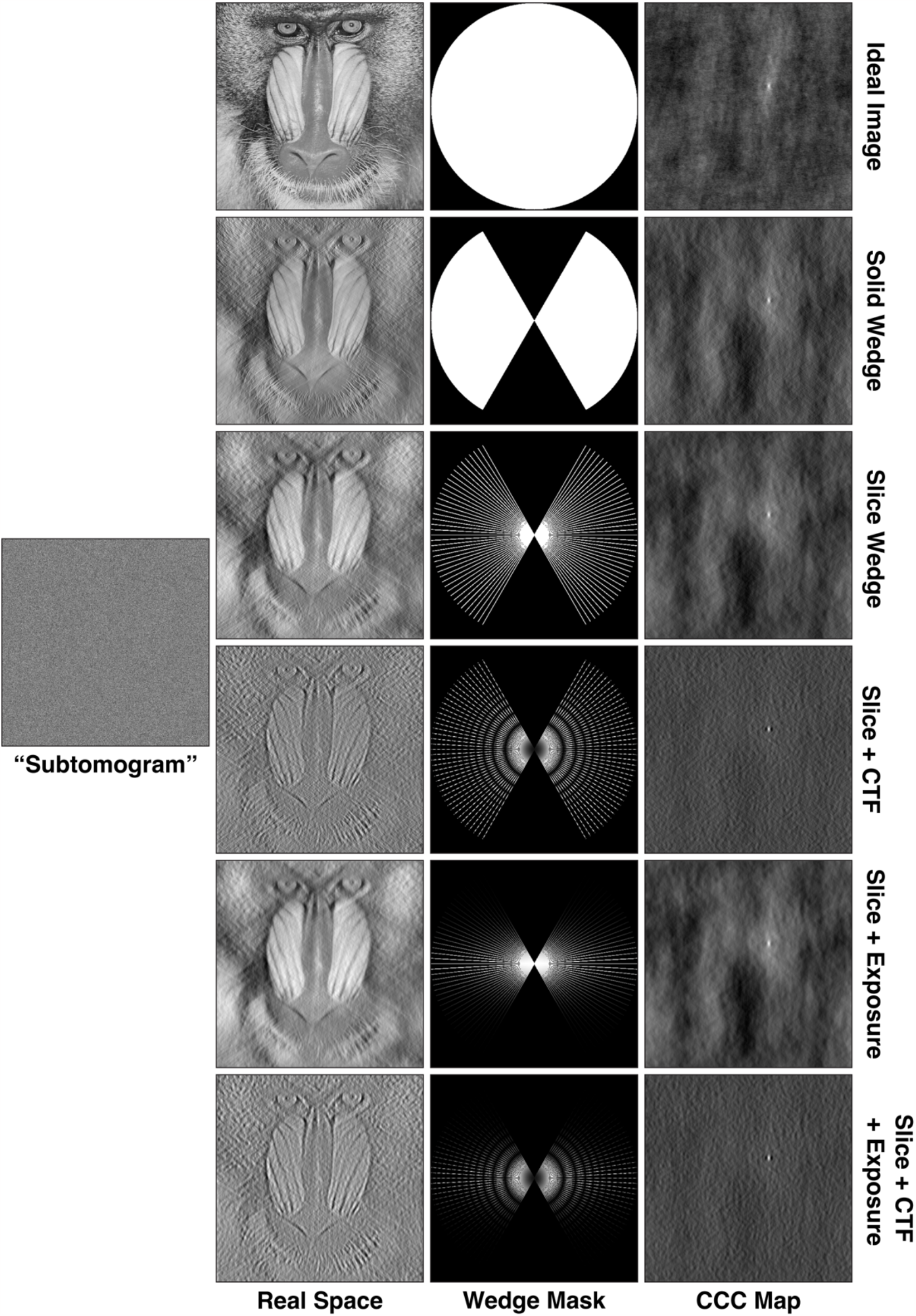
Simulated images with various missing wedge filters and corresponding CCC maps. The ideal mandrill test image has no filters; the “subtomogram” image is the ideal image shifted towards the upper right corner, with a slice wedge, CTF, and exposure filter applied, as well as Gaussian noise at a signal-to-noise ratio of 0.05. The remaining real space images have the noted missing wedge filters applied while the corresponding CCC maps are calculated between the real space images and the simulated subtomogram.

Amplitude modulations in Fourier space cause signal delocalization in real-space, which often leads to high levels of background in the CCC-space. While CTF correction ensures that amplitude modulations are positive, the sinusoidal modulation is always present. By properly accounting for these amplitude modulations in the wedge mask, STOPGAP effectively matches the delocalization between the reference and the tomographic data, resulting in sharp CCC peaks with minimal background noise. To calculate CCCs, STOPGAP first filters templates or references by applying a slice wedge mask with CTF and exposure filter modulations and correlates it to the tomogram or subtomogram using a 3D version of the fast local correlation function (FLCF) (Roseman, 2003; Castaño-Díez *et al*., 2012). The benefits of STOPGAP’s missing wedge model for different STA tasks is shown below.

### Template Matching

Template matching is a reference-based approach that uses a predetermined reference map, i.e. a “template”, for identifying target particles in tomographic data (Fig 2) (Frangakis *et al*., 2002). Templates can either be EM-density maps determined using SPA or STA, or simulated density maps generated from atomic models. During template matching, the template is iteratively rotated through a set of orientations that typically cover all of orientation space with a given angular step size (Fig 2, 3). At each iteration, CCCs are calculated between the template and the tomogram of interest; high valued voxels in the CCC map indicate a potential match for the template in that orientation and position. In the first orientation, the CCC map is stored as a cumulative score map, along with a corresponding orientation map, which stores the template orientation at each tomographic position. While iterating through each orientation, new CCC maps are compared with the score map; voxels with the highest values are stored in the score maps and the corresponding orientations are updated in the orientation map. After all orientations are sampled, the final score map can then be thresholded and peak positions and their corresponding orientations are taken as potential particles.

**Figure 2:**
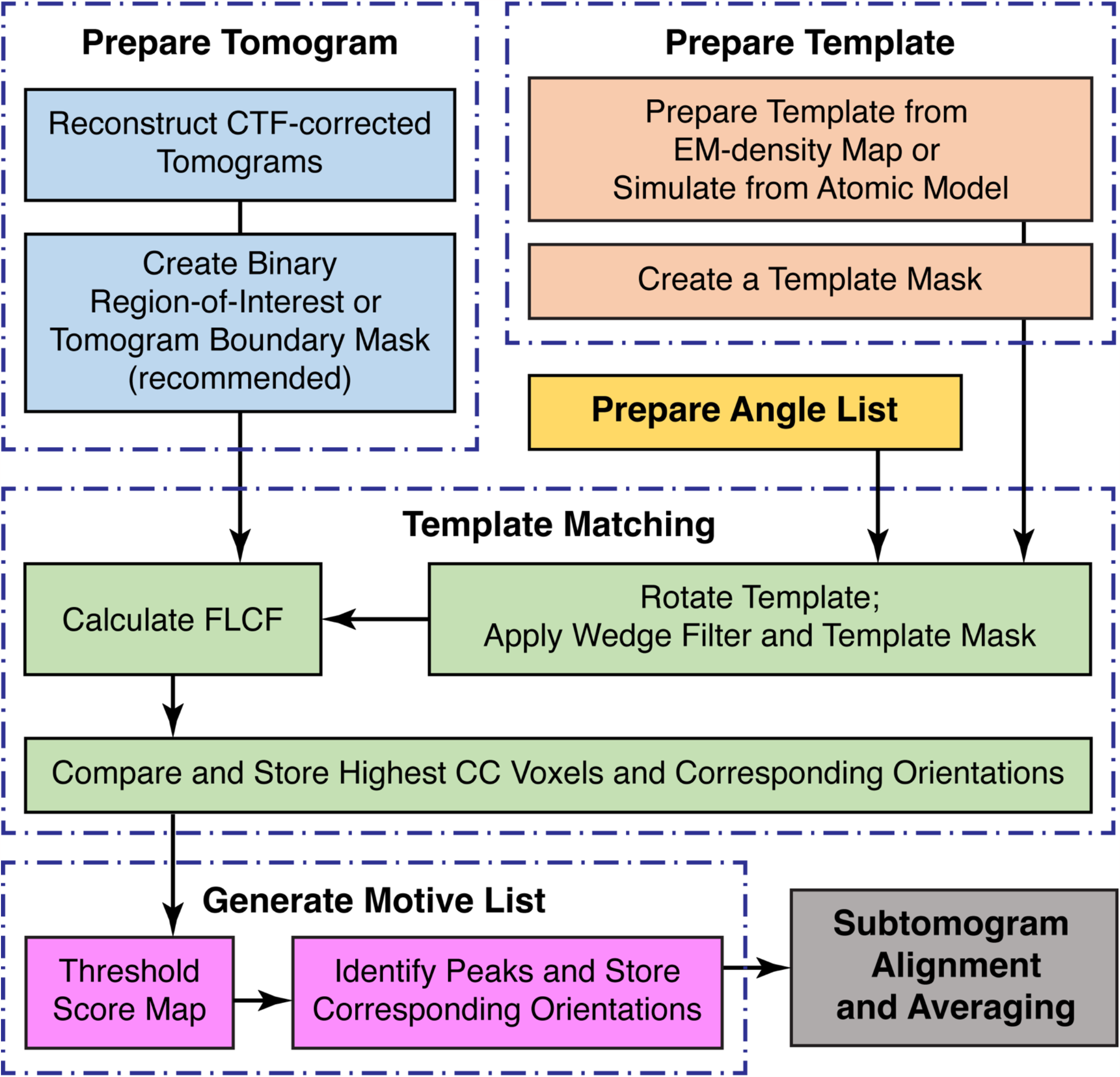
Workflow diagram for template matching. The top row outlines the preparation of the various inputs. The middle row outlines the iterative template matching process. The bottom row outlines the steps for identifying true peaks in the score maps and how they feed into the next steps of the subtomogram averaging workflow.

**Figure 3:**
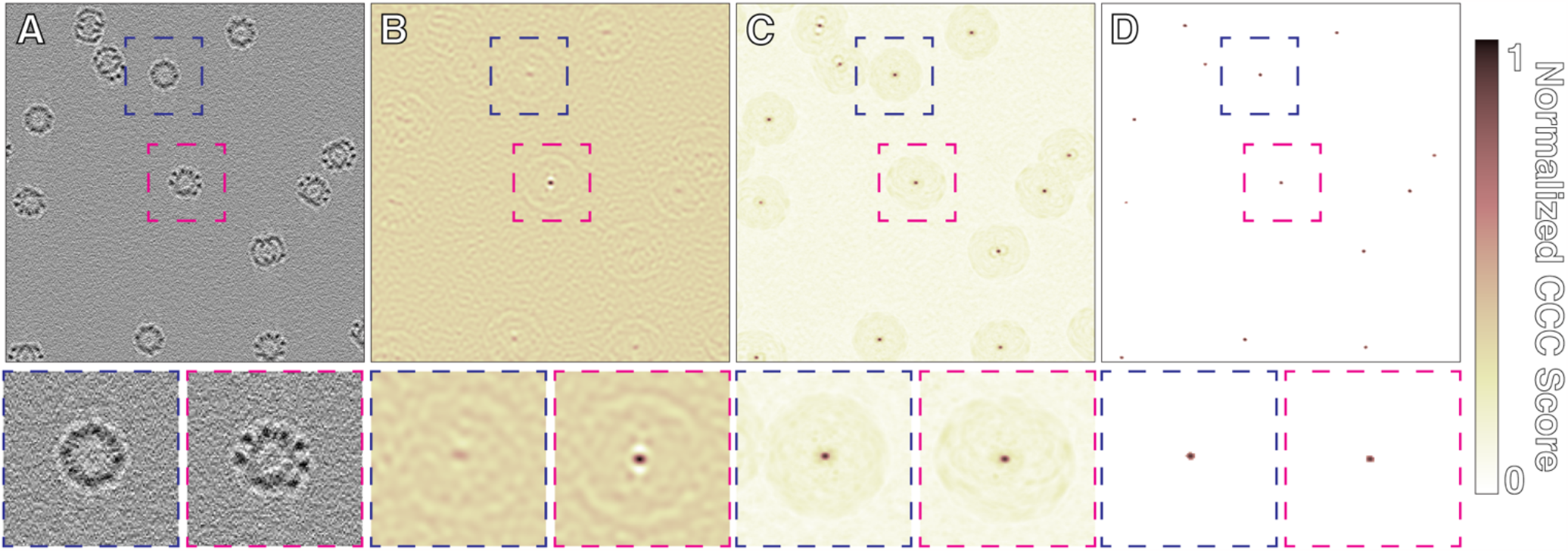
Simulated template matching of thermosomes. A) Slice through simulated tomogram containing randomly placed and oriented thermosomes (PDB: 3J1B). B) Score map from matching with a single orientation. The peak boxed in magenta shows an ideal match with a strong CCC peak; the peak boxed in blue does not match well and has a correspondingly small peak. C) Final score map after full orientational search. D) Score map from C), thresholded to remove background so that only true peaks remain.

Ideally, true positives in the score maps should appear as sharp peaks (Fig 3). However, in many packages, template matching results tend to show a high level of background correlation with CCC value distributions that resemble the input maps (Fig 4A, B). This includes high CCC values for dense objects such as ice contamination or distinct features such as membrane bilayers. By accounting for amplitude modulations, STOPGAP’s missing wedge filter produces CCC peak profiles that are nearly ideal, minimizing noise and false positives (Fig 3 C; 4 C). While sharper peaks have improved signal strength, background noise often takes on a speckled appearance that can make sharp positive peaks difficult to distinguish; this can be a problem when determining the appropriate CCC value to threshold by. To aid with visual analysis, we developed a noise correlation approach where a phase-randomized version of the template is also used for matching. The resultant noise correlation map represents non-specific correlation related to weak structural similarities in the tomogram or template mask-related correlations. The noise correlation map is subtracted from the score map to provide a noise-flattened score map with more pronounced peaks (Fig 4 C,D). For more specifics on the parameters that affect the quality of template matching, a rigorous study on this has been performed using STOPGAP by Cruz-León and colleagues (Cruz-Leon *et al*., 2023).

**Figure 4:**
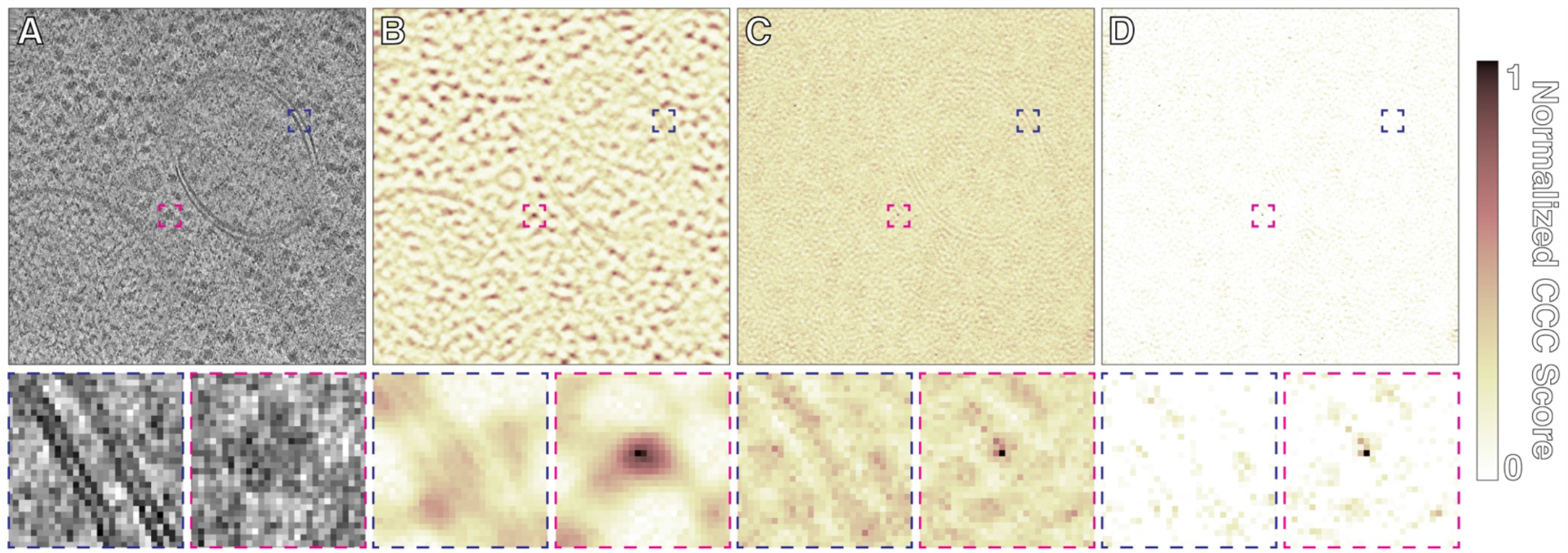
Comparison of template matching in PyTOM and STOPGAP. A) Slice through a tomogram taken from *S. cerevisiae* lamella (*EMPIAR: 11658*). Ribosome template matching with a 15° angular step using B) PyTOM, C) STOPGAP, and D) STOPGAP with noise correlation. Blue boxes indicate CCC scores around a membrane bilayer while magenta boxes indicate scores around a true positive. Colormap shows normalized CCC score, where 1 denotes the value at the CCC peak and 0 the lowest value in the map; score maps in B), C), and D) have not been thresholded.

### Subtomogram Alignment and Averaging

Though we have defined STA broadly as the processing steps that go from tomographic reconstruction up to model building, STA often refers to the determination of higher-resolution structures by aligning and averaging subtomograms. Algorithmically, this can be thought of as two steps: the alignment of a reference to a subtomogram to determine the orientation of that particle; and the generation of a new reference by averaging subtomograms rotated to their determined orientations. Iterating the STA process enables the refinement of subtomogram orientations, as the subtomograms are compared to improved references from the prior iteration.

Subtomogram alignment in STOPGAP is performed in largely the same way as most other real-space packages. A reference map is rotated through a set of orientations, which typically represent a local search in orientational space. At each orientation, the CCC is calculated between the rotated reference and a subtomogram (Roseman, 2003). The maximum value in the CCC map (score), indicates how well the rotated reference matches the subtomogram, while the location of the maximum value in the map provides the Cartesian offset (shift), between the reference and subtomogram. This effectively makes correlation-based STA a 3D rotational search rather than a 6D rotational and translational search, significantly reducing the number of computations. After all orientations are scored, the orientation with the highest score, and its associated shift, are taken as the correct subtomogram alignment.

We performed subtomogram averaging using the 5 tomogram subset of HIV s-CANC particles (EMPIAR-10164) with 3D CTF-corrected tomograms, resulting in a 3.5 Å structure (Fig 5). The most directly comparable published structure is the 3.9 Å structure determined using the AV3 package (Turoňová *et al*., 2017). As with template matching, STOPGAP’s missing wedge filter produces sharper CCC peaks that improve the precision of subtomogram alignment, resulting in higher-resolution averages. Although higher-resolution structures have been determined from this dataset using emClarity, M, and RELION 4 (Himes & Zhang, 2018; Ni *et al*., 2022; Tegunov *et al*., 2021; Zivanov *et al*., 2022), these structures used tilt-series refinement approaches not used here. While tilt-series refinement is not currently implemented in STOPGAP, users have already successfully refined STOPGAP alignments using WARP/M or RELION4 (Khavnekar *et al*., 2022; Rangan *et al*., 2023; Schiøtz *et al*., 2023).

**Figure 5:**
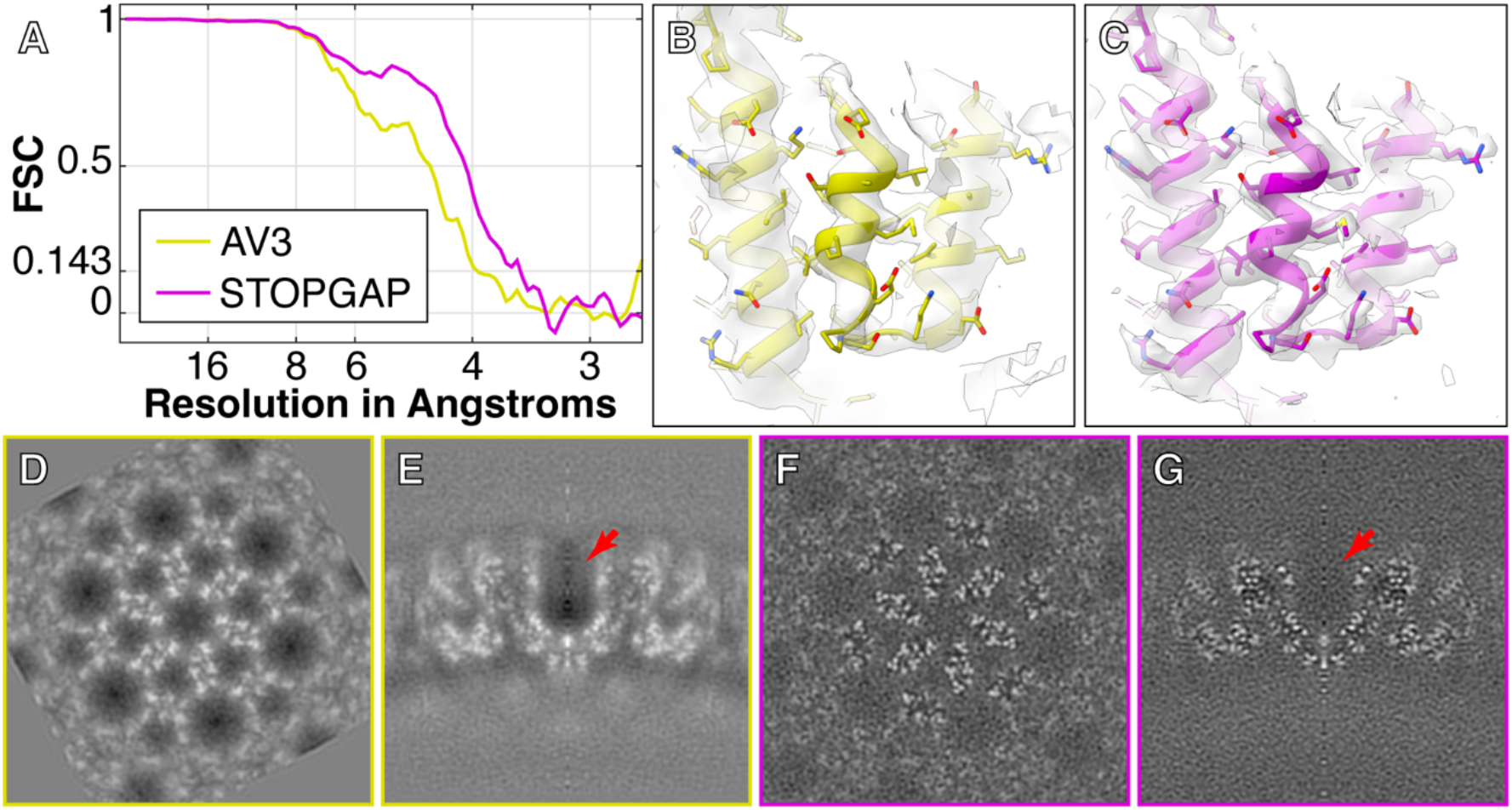
Comparison of HIV s-CANC structure determined by AV3 and STOPGAP. A) Fourier shell correlation (FSC) plots of HIV s-CANC structures determined from the 5 tomogram subset of EMPIAR-10164 by AV3 (Turoňová *et al*., 2017) and STOPGAP. Both used novaCTF (Turoňová *et al*., 2017) for 3D-CTF corrected tomogram reconstruction with no tilt-series refinement. B) and C) are example isosurface representations of the AV3 and STOPGAP EM-density maps, respectively. An atomic model (EMD-3782) is rigid-body fitted into both maps. D-G) XY and XZ orthographic slices through the EM-density maps determined by AV3 and STOPGAP, respectively. Red arrow indicates the high negative density region in the AV3 map that is not present in the STOPGAP map.

In addition to improvements to the estimated resolution, STOPGAP’s missing wedge filter improves the quality and interpretability of averaged maps (Fig 5 B-G). This is because after each subtomogram is rotated, shifted, and summed in real space, Fourier space normalization is required to account for anisotropic sampling. This basically accounts for how the missing wedges from each subtomogram combine to fill Fourier space, which is virtually always anisotropic due to incomplete angular sampling in the dataset. Fourier normalization is performed by rotating and summing the missing wedge filter of each subtomogram, producing a map that tallies the per-voxel sampling in Fourier space. By taking into account amplitude modulations, STOPGAP’s missing wedge filter produces maps with improved normalization. This is illustrated particularly well in subtomogram averages of HIV s-CANC, where normalization with a binary wedge mask produces strong “halos” of negative densities around protein densities (Fig 5 D,E), a phenomenon that has also been noted elsewhere (Sanchez *et al*., 2020). In the STOPGAP average, these areas take on a more ideal appearance (Fig 5 F,G), with grey values around protein densities similar to the surrounding solvent.

### Subtomogram Classification

Classification in STOPGAP is performed using multi-reference alignment (MRA), where each subtomogram is aligned against a set of references, i.e. classes. This effectively makes alignment a 4D search problem, with 3 rotational dimensions and the fourth dimension representing class. References can be from predetermined structures or generated *de novo* from the dataset. Given this, we view MRA into two tasks: the initial generation of *de novo* references, typically from a subset of the dataset; and the alignment of the full dataset against multiple references. In both cases, a key concern is particles becoming trapped in local minimum of orientation space. This can be caused by a number of interrelated issues including initialization bias, where a subtomogram “finds itself” in the reference it contributes to, or premature convergence, where subtomograms stop moving between classes prior to the divergence of structural features in each class.

To overcome these problems, STOPGAP has a number of stochastic methods built into its subtomogram alignment algorithm that can be used during MRA; these are described in detail in the methods section. Briefly, a simulated annealing (SA) algorithm which allows a suboptimal alignment to be accepted with a given probability; this probability is decreased during each iteration of the annealing run. SA encourages more movement of subtomograms between classes during initial phases of MRA. After SA, a stochastic hill climbing (SHC) algorithm is used where the previous orientation and class are scored first, and the order of the remaining orientation and class parameters are randomized. The first better scoring orientation and class is immediately accepted, which allows subtomograms to move more during initial iterations but less as the classes take on distinct features. An additional feature of these stochastic methods is that produce different results when repeating the alignment with the same parameters. We can then use consistency of classification as a rough measure of classification accuracy and precision.

To demonstrate the robust multi-reference classification approach in STOPGAP, we processed *S. cerevisiae* ribosomes from *EMPIAR-11658* (Rangan *et al*., 2023). Using the whole 231 tomogram dataset, we identified ∼240k particles and were able to resolve major structural states of the eukaryotic translation cycle (Fig 6).

**Figure 6:**
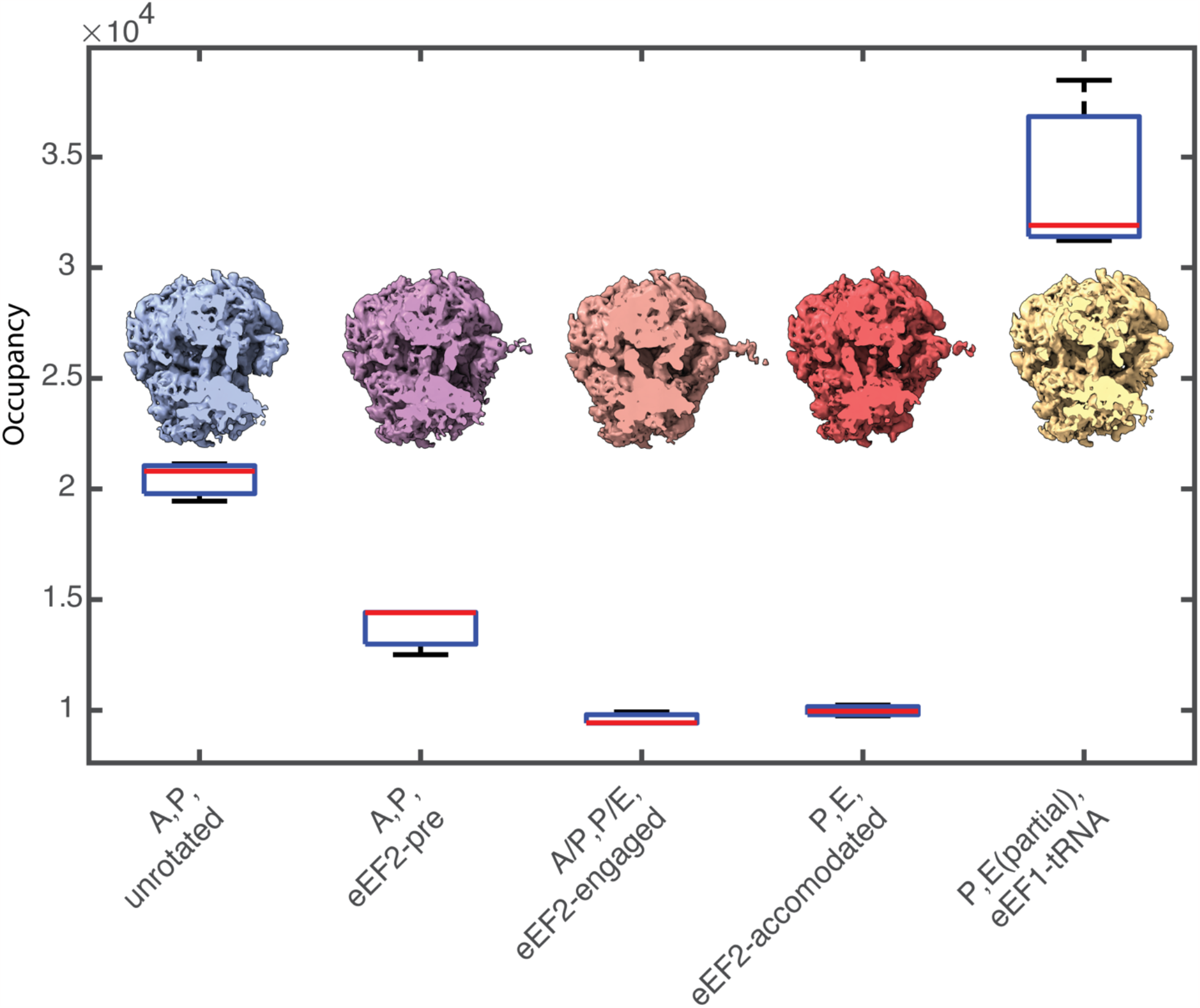
Classification of the *S. cerevisiae* 80S ribosomal translation cycle using MRA. The box plot represents occupancies for each class across three replicates. Red line indicates the sample median, while bottom and top of each box are the 25th and 75th percentiles of the sample, respectively. Whiskers, show the farthest observations beyond the 25^th^ and 75^th^ percentiles. Corresponding density maps for each state after the consensus class assignment are shown with a slice through the tRNA channel. For the ‘P,E(partial),eEF1-tRNA’ state, heterogeneous occupancy of the E-site produces a larger variance across replicates.

Initial ribosome positions and orientations were determined using template matching on 8x binned tomograms, resulting in approximately ∼240K particles. These were then iteratively aligned at 8x and 4x using a mask shaped to contours of the full ribosome density. Particle scores were distributed bimodally; the ∼100K particles in the higher-scoring distribution were selected for further processing. These particles were further aligned at 2x binning, first using a full ribosome mask, followed by alignment using a mask focused on the large subunit. The resulting orientations were used as a starting point for a multireference alignment. Starting references were generated *de novo* by randomly assigning 20% of the dataset to ten classes. To classify the different tRNA states, MRA was performed with a mask focusing on the tRNA channel. The first 10 iterations of MRA were performed using SA, followed by MRA with SHC and without SA until the convergence, i.e. when less than 1% of subtomograms changed classes between iterations. We performed three independent replicates using random *de novo* references and the same MRA parameters. Subtomograms that segregated into the same classes two out of three times were deemed consistent and kept while other particles were deemed unstable and discarded. Final classes were assigned by visually curating the final volumes and merging identical states (Si Fig 1). This resulted in 5 unique states (Fig 6), namely, ‘A, P, unrotated’, ‘A, P, eEF2-pre’, ‘A/P, P/E, eEF2-engaged’, ‘E, P, eEF2-accomodated’, and ‘E(partial), P, t-RNA-eEF1’ (Milicevic *et al*., 2023). Class occupancies between replicates were similar (Fig 6) indicating the reproducible separation of stable classes while also providing a metric for quantitating the class assignments. In this instance, we defined class consensus as a subtomogram assignment to the same class two out of three times. The number of replicates and stringency for consensus are ultimately defined by the user, depending on their specific needs (Erdmann *et al*., 2021). Final averages were generated using a consensus subtomogram assignment for each of the three classes.

## Discussion

STOPGAP is a MATLAB-based open-source package for STA. At its core is a missing wedge model designed to account for the various types sampling and modulation effects that occur during tomographic data collection and reconstruction. This missing wedge model improves the performance of the 3D CCC, which subsequently improves the performance of template matching and subtomogram alignment. This missing wedge model also improves the Fourier-space normalization function used in STA, providing improved EM-density maps. For template matching, we also developed a noise-correlation approach to reduce background correlations in template matching score maps and enhance the appearance of true peaks. To facilitate MRA, we introduced SA and SHC algorithms to our subtomogram alignment procedure to help overcome convergence on local minima as well as provide metrics for assessing the reproducibility and reliability of subtomogram classifications. Such assessments provide more quantitative values for class occupancies, which is of particular importance for cellular cryo-ET as quantitatively assignment of conformational states is important for characterizing biological states.

STOPGAP primarily focuses on the aspects of STA that use real-space correlation approaches with the aim of providing users fine-grained control in how their data is processed. To this end, an extensive range of parameters are open to the user for fine tuning, though most come with presets that are suitable to a wide range of problems. STOPGAP is not intended to be a complete and comprehensive package for all aspects of STA, but instead aims to provide a modular set of tools for carrying out specific image processing tasks. Given the rich information content of cryo-ET data, we believe that there is no single pipeline that can answer every biological question; STOPGAP’s task-oriented approach allows users to tailor pipelines suited to their particular biological questions. Given that STOPGAP is unlikely to be the optimal solution to all cryo-ET problems, it is also aimed to be complementary with other cryo-ET packages, enabling users to build comprehensive STA workflows that meet the demands of their specific projects. A number of studies have already been performed using STOPGAP as a component of the processing workflow alongside packages such as Warp, M, RELION, and novaSTA (Hoffmann *et al*., 2022; Xing *et al*., 2023; Rangan *et al*., 2023; Khavnekar *et al*., 2022, 2023; Schiøtz *et al*., 2023; Turoňová *et al*., 2020; Erdmann *et al*., 2021; Lacey *et al*., 2023). Overall, we believe that STOPGAP provides a powerful set of image processing algorithms and tools that are complementary to others in the community and hope that the descriptions of our algorithms will useful for further community-driven development. STOPGAP is available at https://github.com/wan-lab-vanderbilt/STOPGAP.

## Methods

### Software Overview

STOPGAP is an open-source package written in MATLAB, consisting of a main STOPGAP executable, which is run as a compiled MATLAB executable, and the “toolbox,” a set of MATLAB functions and scripts. The STOPGAP executable is supplied pre-compiled, but can also be compiled by end users. Example bash scripts for running STOPGAP in parallel using the message passing interface (MPI) or SLURM Workload Manager are provided, though these will likely need to be edited to match specific cluster configurations.

In addition to the main executable, a “parser” executable is also supplied, which checks for conflicting parameters and generates properly formatted parameter files for the various STOPGAP tasks. STOPGAP tasks include subtomogram extraction, template matching, and subtomogram alignment. MRA-based classification is not a separate task but instead a user-implemented workflow using the subtomogram alignment task.

### Missing Wedge Model

The missing wedge is modelled as a series of Fourier slices, each of which can have additional amplitude modulations corresponding to the local CTF of each subtomogram and the exposure filter applied during tomogram reconstruction. For template matching, the CTF filter is calculated as a single global filter consistent throughout the tomogram. Local per-particle CTF filters are used for subtomogram alignment, averaging, and classification. Local CTF filters are calculated by first assuming that the defocus estimated from a tilt-series represents the defocus at the center of mass of the specimen (Turoňová *et al*., 2017). The center of mass of the tomogram is estimated using the particle positions in the “motivelist”, which stores the alignment parameters of each subtomogram. For each Fourier slice, the local defocus of the subtomogram is calculated using the estimated defocus, the center of mass, distance of from the tilt axis in each tilt projection.

### Constrained Cross Correlation

CCC’s are calculated using a 3D version of the FLCF (Roseman, 2003; Castaño-Díez *et al*., 2012). The FLCF is a real-space correlation function that normalizes the reference map (i.e. reference or template) and the search map (i.e. tomogram or subtomogram) with respect to the mask applied to the reference. Prior to calculation of the FLCF, reference maps are filtered with the amplitude-modulated missing wedge filter while search maps are filtered with a binary slice wedge masks to remove any reconstruction- or cropping-related Fourier space artifacts.

### Template Matching

During template matching, each tomogram is split into subvolumes, termed “tiles”, for parallel processing. For each tile, the template is rotated through a set of orientations defined in the “angle list” and a CCC is calculated from the rotated template and tile. Highest scoring voxels for each orientation are accumulated into a “score map” while associated orientations are stored in the “orientation map”. Motivelists are then generated by first thresholding score maps to a user-defined level, followed by a peak finding algorithm that searches for the highest scoring voxels with a user-supplied inter-peak distance related to the particle dimensions. Particle positions are taken from the peak positions in the score map while particle orientations are taken from the same positions in the orientation map.

STOPGAP template matching also includes an additional noise-correlation approach, similar to that used in calculating Fourier shell correlations (Chen *et al*., 2013). In this approach, a noise volume is first generated by randomizing the phases of the template. This noise template is used for matching alongside the actual template, resulting a noise score map that is used to reweight the main score map. Reweighting has the effect of shifting the distribution of the CCC values downwards. Negative correlations in the reweighted score map are then set to zero, which effectively flattens noise, allowing for better visual analysis for score thresholding.

### Subtomogram Extraction

Subtomogram extraction is performed by cropping subvolumes from each tomogram according to the positions stored in the motivelist. Positions in the motivelist refer to the center of each subtomogram. The default file format for subtomograms is the .mrc format, while .em format is also supported; subtomograms saved as .mrc files can be stored as 8-16- or 32-bit data.

### Subtomogram Alignment and Averaging

Subtomogram alignment revolves around the parameters stored in the motivelist, primarily the CCC score, shift, and rotation values from prior iterations. Input for an alignment round include bandpass filter settings, angular increments which define the granularity of the search space, and the angular iterations, which define the size of the search space. The angular increments and iterations are used to calculate a list of search orientations with respect to the prior determined orientation. At each orientation the CCC is calculated; the peak value is taken as the CCC score while the vector from the center of the CCC map to the peak is taken as the shift vector. A cross-correlation mask can be applied to limit the shifts allowed. As alignment progresses, the angular increments are decreased to finer angles to align higher resolution features. After alignment is completed, a new motivelist with updated CCC scores, shifts, and rotations is produced. This new motivelist is then used to average new references.

Subtomogram alignment and averaging is always performed as “halfsets” where the motivelist is split into two and aligned and averaged independently. For so-called “gold-standard” alignment (Scheres & Chen, 2012), where halfsets are aligned and averaged completely independently, the halfset of each subtomogram must be defined in the motivelist. If halfsets are not defined, STOPGAP will randomly split the dataset in half during each iteration.

### Classification by MRA

In standard single-reference alignment, STOPGAP is essentially performing a 3D search of orientation space, with the 3D translation implicitly determined as the peak position in the CCC map. MRA is formulated as a 4D search, with the fourth dimension denoting the references or classes. As such, the set of search orientations not only contains the three Euler angles, but also each reference. Standard subtomogram alignment takes the form of an expectation maximization or hill climbing algorithm, where the complete set of search orientations is scored and the maximum scoring orientation is accepted. STOPGAP alignment includes a SHC approach inspired by similar SPA approaches (Reboul *et al*., 2016), where the previous best orientation is scored first while the rest of the orientations are scored in random order. The orientation that scores better than the previous orientation is immediately accepted and STOPGAP moves on to the next subtomogram. SHC ensures that better scoring orientations are found at each iteration, but prevents being trapped in local optima by not searching for the “best” orientation. An additional simulated annealing (SA) method is also implemented on top of the SHC approach, which allows for the acceptance of worse scoring orientations. In SA, after scoring the previous best orientation, a worse scoring orientation can be accepted with a given input probability; the annealing occurs by decreasing this probability to zero over subsequent iterations.

*De novo* reference generation refers to the process of generating multiple references from the dataset without *a priori* knowledge of the class of each subtomogram. This is typically performed on a subset of the data by first deciding on the number of classes to be produced, then randomly splitting the dataset into even bins for each class followed by MRA. To minimize the potential for initialization bias, STOPGAP can initialize *de novo* references by asking the user for the number of desired classes and the number of subtomograms per class. It then randomly selects subtomograms to populate each class in a non-exclusive manner. This is different from evenly splitting the dataset as a subtomogram may appear in multiple classes or none during initialization, reducing the potential and impact of the attractor problem (Sorzano *et al*., 2010).

Following *de novo* reference generation, MRA is then performed on the initial references to refine them into divergent structures. MRA is first performed using SA, which has the effect of increasing particle movement between classes, followed by SHC until convergence. STOPGAP’s stochastic methods have an additional benefit that MRA can be performed with the same parameter set to produce different results, which we use as a way of judging the reproducibility of our classification. Repeated MRA runs that produce classes with consistent sizes suggests stable separation of conformations as well as provides an estimate of the error of class occupancies. Likewise, subtomograms that repeatedly find the same class are likely to be true members of that class while those that converge on to different classes at each MRA run suggests bad particles that can be removed from the dataset.

## Acknowledgements

This work was supported by U.S. National Institutes of Health grant DP2GM146321 (to WW). WW is a Pew Scholar in the Biomedical Sciences, supported by the Pew Charitable Trusts. This work was conducted in part using the resources of the Advanced Computing Center for Research and Education at Vanderbilt University, Nashville, TN. Some of this work was performed at the Max Planck Institute of Biochemistry with support from Wolfgang Baumeister, Juergen Plitzko, and John Briggs for resources and infrastructure. We would also like to tank Max Planck Computing and Data Facility (MPCDF) for the computational infrastructure. SK would like to thank the Max Planck Society and the International Max Planck Research School for graduate school funding.

---

**Supplementary Figure 1:**
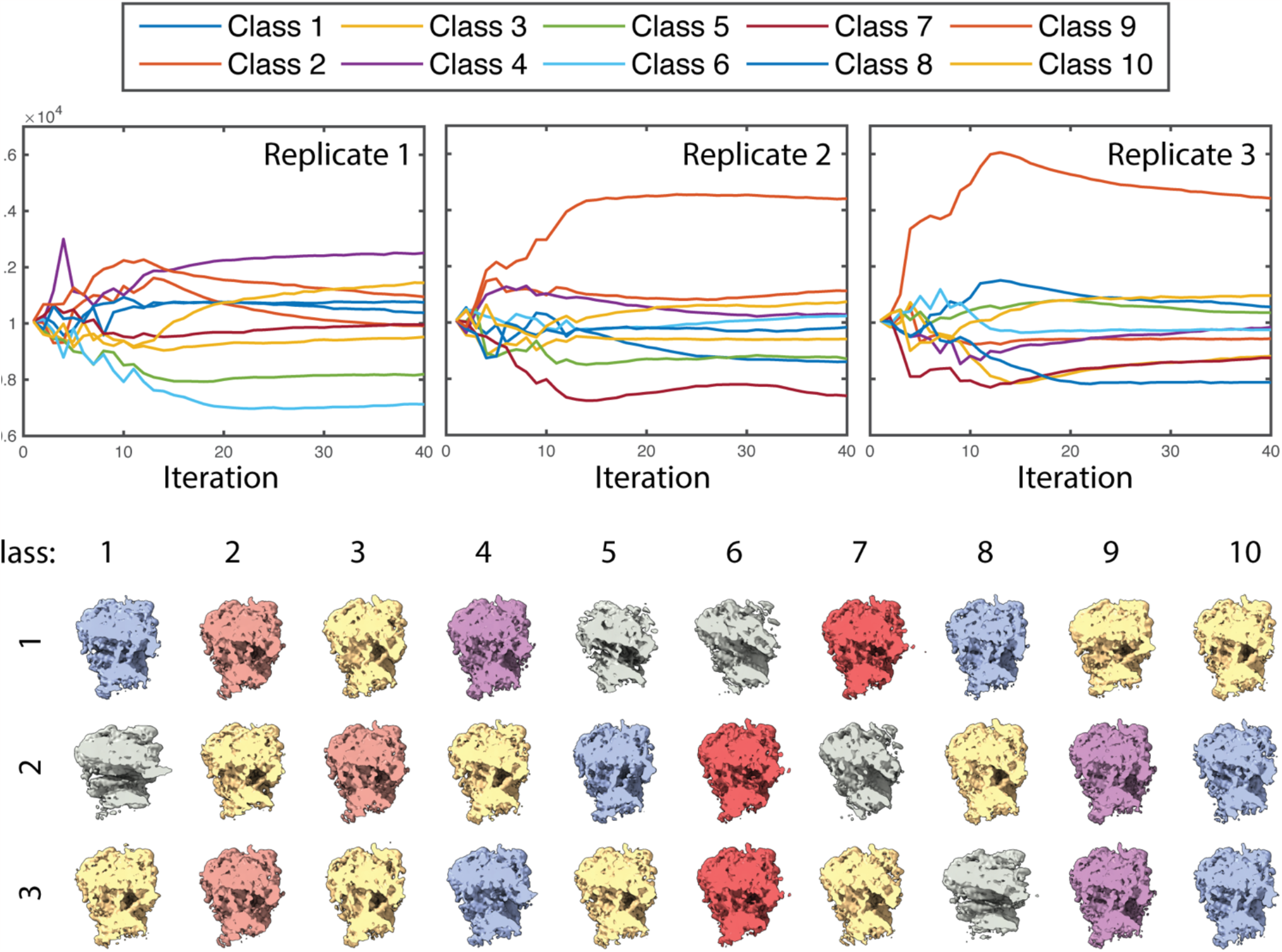
Classification of the *S. cerevisiae* 80S ribosome over three replicates. A) Occupancy of classes over each iteration for each of the three iterations. Class numbers are prior to final curation and are arbitrary between replicates. B) Final density maps after MRA classification with 10 classes across three replicates. Density maps are colored by the visually curated and assigned states depicted in figure 6.

